# Whole transcriptome sequencing and biomineralization gene architecture associated with cultured pearl quality traits in the pearl oyster, *Pinctada margaritifera*

**DOI:** 10.1101/421545

**Authors:** J. Le Luyer, P. Auffret, V. Quillien, N. Leclerc, C. Reisser, J. Vidal-Dupiol, C.-L. Ky

## Abstract

**Background:** Cultured pearls are unique gems produced by living organisms, mainly molluscs of the *Pinctada* genus, through the biomineralization properties of pearl sac tissue. Improvement of *P. margaritifera* pearl quality is one of the biggest challenges that Polynesian research has faced to date. To achieve this goal, a better understanding of the complex mechanisms related to nacre and pearl formation is essential and can now be approached through the use of massive parallel sequencing technologies. The aim of this study was to use RNA-seq to compare whole transcriptome expression of pearl sacs that had producing pearls with high and low quality. For this purpose, a comprehensive reference transcriptome of *P. margaritifera* was built based on multi-tissue sampling (mantle, gonad, whole animal), including different living stages (juvenile, adults) and phenotypes (colour morphotypes, sex).

**Results:** Strikingly, few genes were found to be up-regulated for high quality pearls (n = 16) compared to the up-regulated genes in low quality pearls (n = 246). Biomineralization genes up-regulated in low quality pearls were specific to prismatic and prism-nacre layers. Alternative splicing was further identified in several key biomineralization genes based on a recent *P. margaritifera* draft genome.

**Conclusion:** This study lifts the veil on the multi-level regulation of biomineralization genes associated with pearl quality determination.

## Background

The mollusc *Pinctada margaritifera* var. *Cumingii* is a species of great economic importance in French Polynesia. The associated pearl industry represents the second most important source of income there, just after tourism. Cultured pearl production requires two distinct animals. A small piece of graft mantle tissue is dissected from a sacrificed donor oyster and inserted with a round bead of nacre (a nucleus, made of mussel shell) into the gonad of a recipient oyster [1, 2]. If the graft is not rejected and the recipient oyster survives the grafting operation, the implanted mantle tissue will grow to completely surround the nucleus and form a “pearl sac”, capable of secreting biomaterial layers (calcite and aragonite) around the nucleus [3]. After 15 to 24 months of culture, pearls are harvested and usually sorted according to six main quality traits: size, shape, colour, surface complexion, lustre and grade [4]. It is estimated, however, that only 5% of the harvested pearls can be classified as grade A, which corresponds to the best quality according to the local regulatory quality standards [5]. Average export price of cultured pearls in French Polynesia has fallen dramatically over the past decade, mainly due to a combination of factors including overproduction; hence quantity has been favoured to the detriment of quality. The improvement of cultured pearl quality is an imperative aspect of a pearl farm’s sustainability and as one of the biggest challenges that research is facing in French Polynesia.

Factors affecting pearl quality have diverse and non-exclusive origins including genetics, environment and/or genotype-by-environment interaction (GEI). This is further complexified in the *Pinctada* transplant model because of the phenotype transmission from the donor oyster to the recipient oyster, their interplay, and their interaction with the environment. One particularity of this animal model is the chimera system attributed to the pearl sac tissue, whose interaction between Genome_Donor_ × Genome_Host_ has a significant effect on pearl quality. A recent study in *P. margaritifera*, based on controlled bi-parental crosses and the F1 generation, demonstrated heritability (*h*^2^ from 0.21 to 0.37) for nacre weight and thickness, pigmentation darkness and colour, surface defects and grade, signifying a donor oyster effect with a genetic basis, although there were also important interaction components [6]. Previous studies reported that location, temperature and food availability [7–9], pearl rotation [10], donor oyster genotype [4, 11–14], age [15], position of graft mantle [16] and contamination during the graft operation and/or graft operator skills [17] are all determinants of final pearl quality and do not necessarily affect similar traits.

Various genomic approaches have been applied in pearl oyster with the objectives of identifying key candidate markers related to pearl quality traits. For instance, Lemer et al. [18] identified a set of genes differentially expressed between two matle colour phenotypes (black *P. margaritifera* phenotypes and full albino individuals), using a suppressive and subtractive hybridization (SSH) method. In silver-lipped oyster (*Pinctada maxima*), genetic association analyses has permitted the identification of QTLs linked to pearl surface complexion and colour as well as genetic associations of regions and markers for pearl size, weight, colour and surface complexion [19, 20]. Transcriptome-wide and proteomic approaches have also been used to characterize the pool of genes expressed during pearl formation and to discriminate markers preferentially associated with nacreous and/or prismatic layers. Studies in the Japanese pearl oyster (*Pinctada fucata*) showed that the genes *msi60* and *aspein* from the mantle tissue were up-regulated in low quality pearls compared to high quality groups, while the expression of the *msi30* gene from the pearl sac tissue was up-regulated in high quality pearls [21, 22]. Finally, a recent study on *P. margaritifera* revealed that *shematrin5 and 9, prismalin* and *aspein* encoding genes were up-regulated in the pearl sacs of individuals producing low pearl surface quality [23]. Studies have been limited to relatively few candidate genes, however, and an overall evaluation of the actors involved in pearl quality remains to be conducted.

The aim of our present study was to identify key genes involved in the regulation of pearl quality through a comparative RNA-seq analysis of pearl sacs producing high and low quality cultured pearls. We constructed a new comprehensive multi-tissue transcriptome assembly, covering several developmental stages, colour morphotypes and tissue origins, which will be useful for further transcriptomic studies in *P. margaritifera*. Furthermore, based on a recently assembled draft genome of *P. margaritifera*, we successfully explored the possibility that alternative gene splicing events are involved in the regulation of biomineralization processes.

## Results

### Transcriptome assembly

The raw transcriptome assembly contained 541,184 transcripts. After filtering for redundancy and functionality, we retained a total of 41,075 transcripts (assembly metrics given in Table 2). Transcriptome completeness evaluation indicated that 90.6 % of the highly conserved single-copy metazoan genes (n = 978) were present in our transcriptome (89.8 % are complete and in a single-copy). Similarly, mean mapping rate reached 66.31 ± 1.72 % with negligible differences in sample condition. Both the transcriptome completeness and satisfactory mapping rate suggest that the several steps of filtering applied did not have major impact on the overall transcriptome. Functional annotation identified a total of 33,532 transcripts (81.5%) with at least one match with deposited sequences (Table 2).

### Differential expression in biomineralization-related genes

We found a total of 262 differentially expressed genes (DEGs), with 246 up- and 16 down-regulated genes in low quality pearls (Table S3). Out of the 262 DEGs, 216 (82.24%) had at least one match with known protein sequences (Table S3). Finally, only 114 of the DEGs (43.5 %) had at least one associated GO term. GO analysis revealed enrichment for some relevant functions involved in pearl formation, including oxidoreductase activity (GO:0016491), peptidase inhibitor activity (GO:0030414), serine-type peptidase activity (GO:0008236), chitin binding (GO:0008061) and copper ion binding (GO:0005507). The GO enrichment analysis is summarized in Figure S1.

We identified several biomineralization genes discriminating high and low quality pearls (Figure 3; Table S3). Most of these genes are characteristic of prismatic and prism-nacre layers [24] and were found up-regulated in low quality pearls. The blue mussel shell protein-like (*BMSP-like*) coding gene is the single biomineralization gene up-regulated in high quality pearls. *BMSP-like* shares strong homology with the *P. fucata pif-177* gene and is notably involved in determining the polymorph of CaCO_3_ [25]. We also identified Gypsy and Jockey-family transposable elements up-regulated in high quality pearls (Table S3). The qPCR analysis shows that the individual relative expression of the four biomineralization-related genes analysed is in accordance with results from pool RNA-seq data (Figure S2). We found the *MP10, shematrin-9* and *aspein* up-regulated in low quality pearls (p-value < 0.001) while no significant difference was observed for *pif-177* (p-value = 0.012).

**Figure 1:**
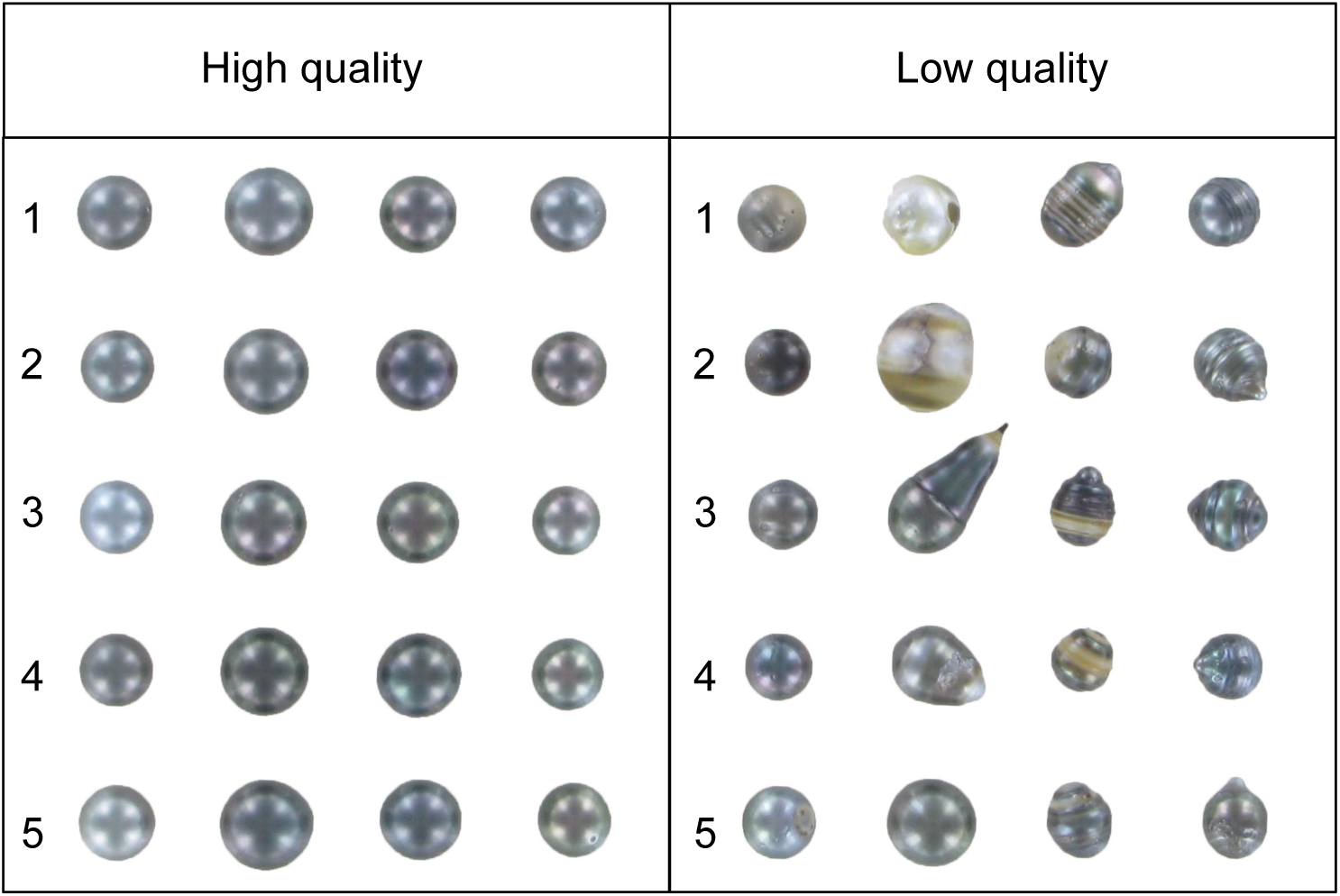
High and low quality cultured pearl samples from *P. margaritifera*. Each row within each condition (high or low quality), represents a specific pool (n = 4 cultured pearls / pool / condition). Numbers represent pools by condition (high or low pearl quality).

**Figure 2:**
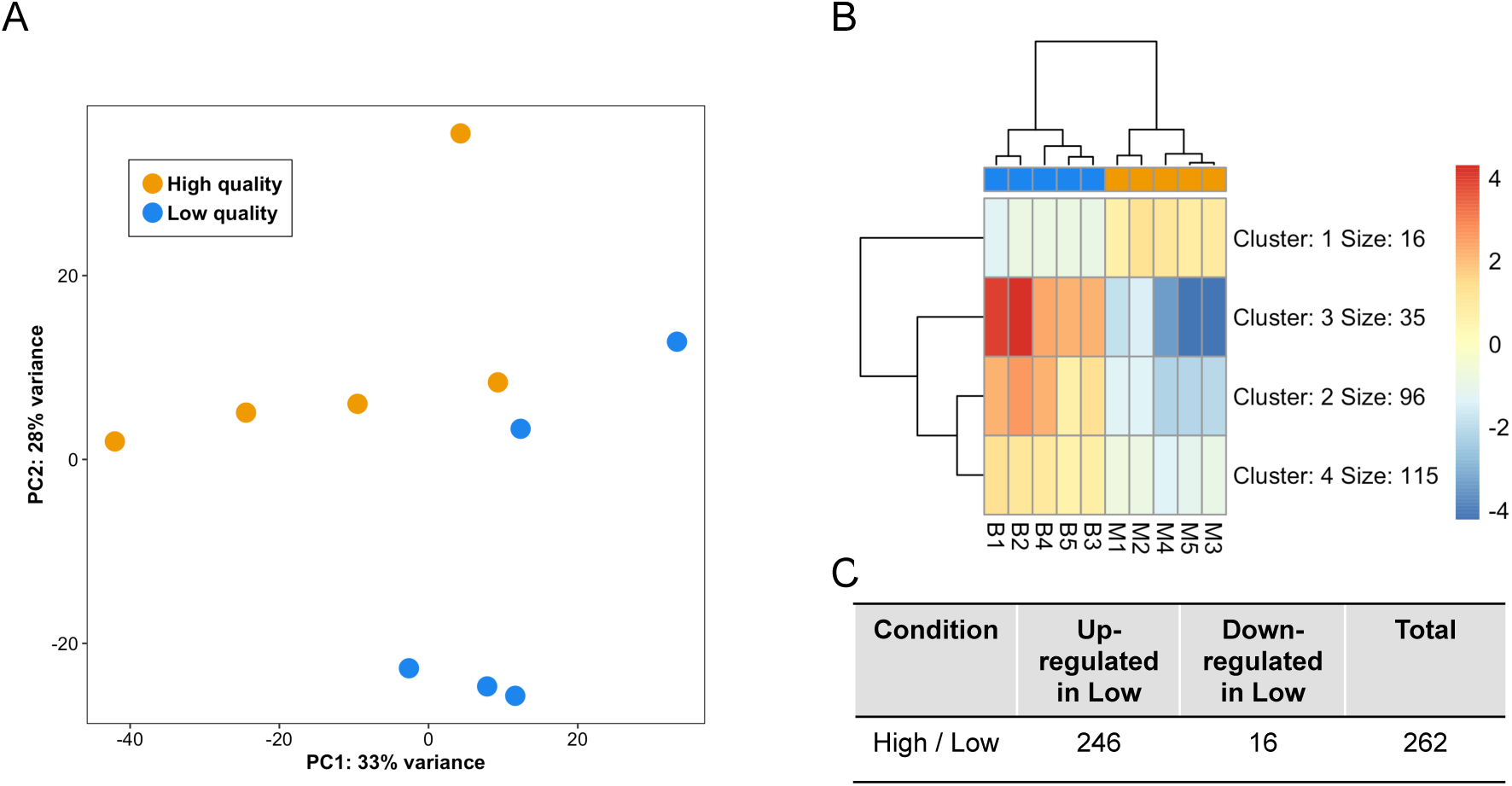
Genes differentially expressed between *P. margaritifera* pearl sacs having produced high and low quality pearls: A) Principal component analysis (PCA); B) Heatmap of differentially expressed genes. LogCPM (+2) were computed based on raw counts normalized for library size reported to the mean gene expression over all individuals; and C) Table showing the number of differentially expressed genes.

**Figure 3:**
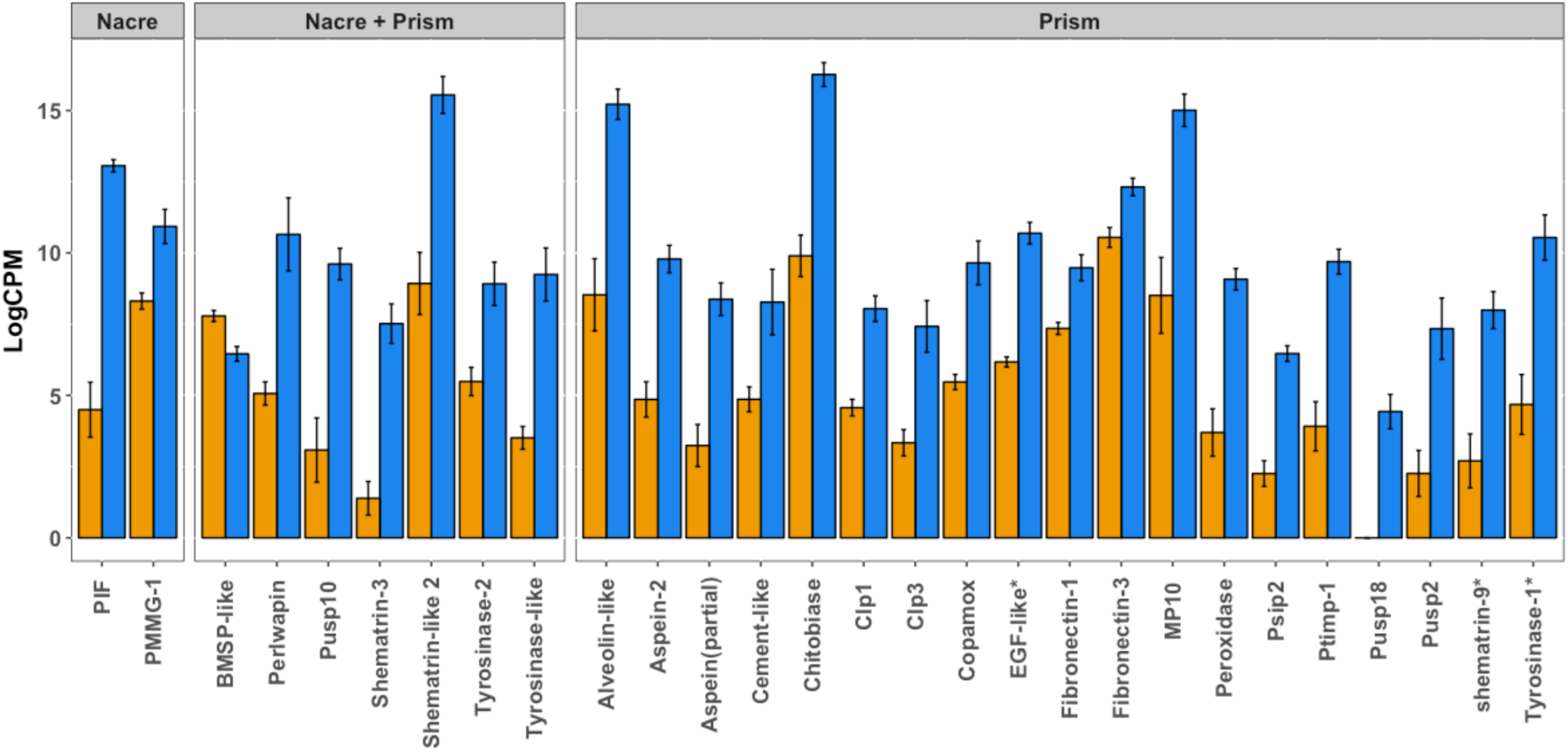
Bar plots of mean expression of biomineralization-related genes in pearl sac of *P. margaritifera*. Only genes with significant differential expression (FDR < 0.01 and |log2FC| > 1) are reported for clarity. Values are expressed as the mean (logCPM + 2) per condition ± standard deviation. Orange = high pearl quality; Blue = low pearl quality. Asterisks indicate genes for which multiple isoforms were reported in the transcriptome assembly. For each of these genes at least one of the isoforms was differentially expressed.

### Different alternative splicing in biomineralization-related genes

We found a total of 28 transcripts showing significant differential splicing (FDR < 0.001) (Figure 4). Several of these genes are known to be involved in biomineralization processes: *pif, aspein, pwap* and *wdf18*. For *aspein*, high quality pearls show lower exon 1 usage. This specific exon overlaps with the promoter domain of the transcript while the exon 2 codes for the D-domain rich in Asp amino acid that gives the aspein protein its crystal binding affinity and its function in regulating crystal growth [21, 26–29]. The GO enrichment analysis identified the peptide biosynthetic process (GO:0043043), amide biosynthetic process (GO:0043604) and translation function (GO:0006412) as enriched for the genes showing different exon usage.

**Figure 4:**
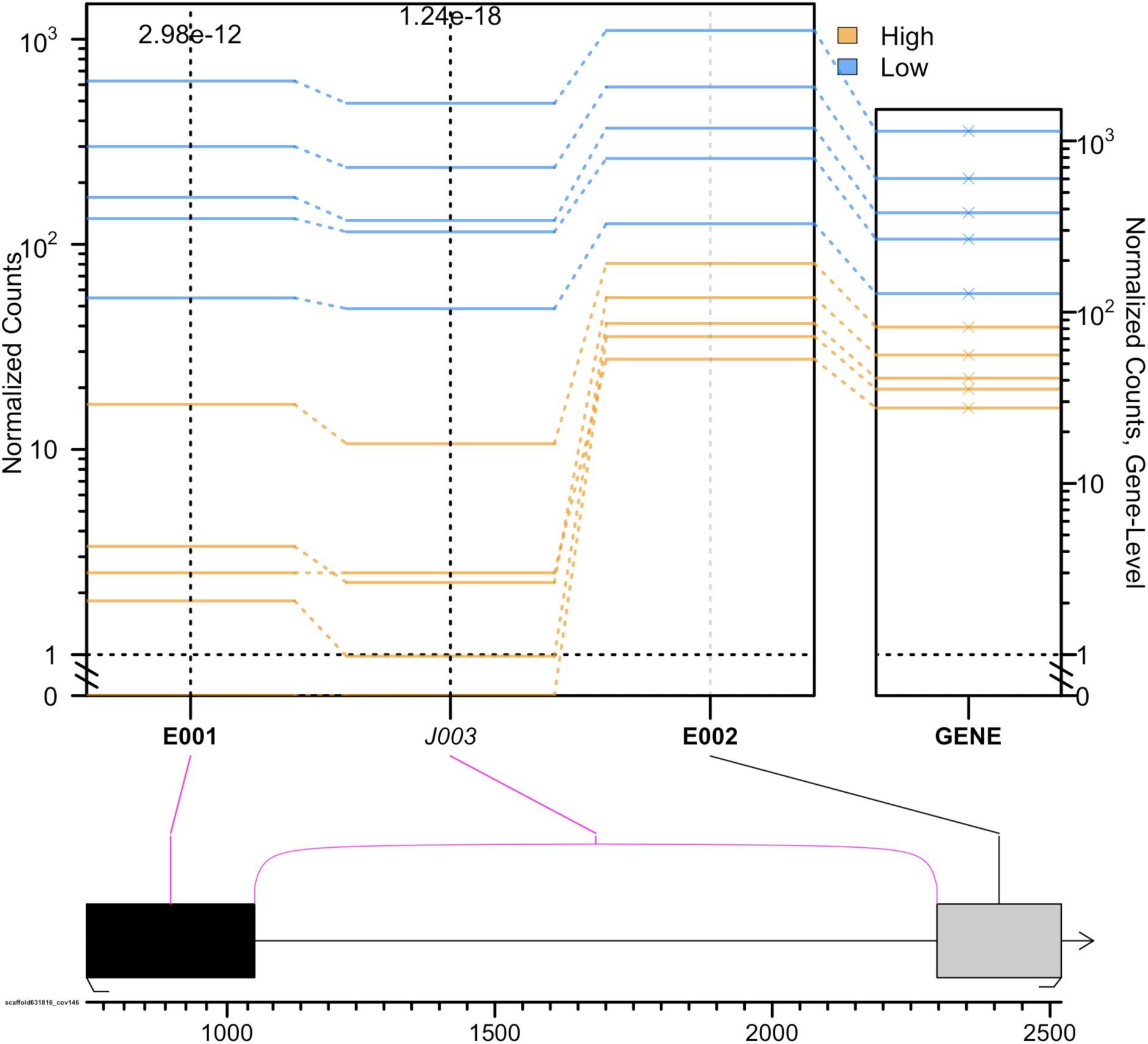
Splicing event visualization for the aspein gene in the pearl sac of *P. margaritifera*. Normalized counts are plotted for each gene section, either exon (E) or junction (J), and each individual; blue = high quality and orange = low quality. Values in the box plot represent p-values (Fisher’s test) for each gene section.

## Discussion

Access to massive parallel sequencing technologies now greatly contributes to the understanding of molecular expression of the phenotype in non-model species. Here, we used a common transcriptomic approach (RNA-seq) to obtain an overview of differences in gene expression and alternative splicing between high and low cultured pearl quality. Recently, RNA-seq has been successfully used to explore genes related to pearl oyster growth and response to environmental stressors (*P. fucata*) [30, 31] and biomineralization (*P. martensii* and *P. penguin*) [32–34]. The use of RNA-seq in *P. margaritifera* for biomineralization related studies has been held back by the relatively limited coverage of previous transcriptome references (Roche 454; [35]) or their reduced tissue representation [36]. Recent work has however completed multi-tissue transcriptome assemblies of four species of pearl oysters including *P. margaritifera*. Yet, unfortunately, sampling did not include pearl sac tissues and to our knowledge assemblies were not made publicly available [37]. The present study provides a comprehensive reference transcriptome for *P. margaritifera*, encompassing whole-body tissue as well as phenotypic variation for a single common tissue (for gonads this was either female or male; for mantle tissue and whole individuals this covered a broad range of shell colour) as well as including individuals of different life stages (juveniles or adults). Even when applying stringent filtering, the satisfactory completeness as well as good individuals mapping rates and high gene content conservation with the closely related species *P. fucata* (78% overlap) all suggested that this new reference transcriptome should prove itself a useful genomic resource for a broad range of future transcriptomic research for *P. margaritifera*.

### Complex responses of biomineralization genes are associated with pearl quality

The pearl sacs producing low quality pearls were characterized by a higher activity of prismatic layer-specific genes. Among the differentially expressed genes, *aspein* and *shematrin-9* were found to be up-regulated in low quality pearl RNA-seq pools, as well as being validated by individual qPCR, which is consistent with previous candidate gene expression work [23]. The present study also provides a novel set of biomarkers involved in the biomineralization process, such as *perlwapin* and *BMSP-like* genes, and supports previous findings showing deep conservation of biomineralization genes within molluscs [38–40].

A considerable number of studies have focused on the identification of genes involved in aragonite and/or calcite formation in pearl oyster species and other bivalves capable of shell mineralisation [16,18,45–51]. Nonetheless, it has proven difficult to extrapolate the role of key actors involved in determining cultured pearl quality across studies. It is reasonable to hypothesize that contrasting results in the expression of biomineralization genes might result from the pleiotropic effect of biomineralization genes and/or major differences in the design of these different studies (genetic background, geographical locations, pearl grades sampled, time of sampling post-graft, environmental factors and tissue-analysed). For instance, differences in response of the *pearlin* gene, a gene commonly used to monitor biomineralization, have been observed in both graft and pearl sac tissues under similar environmental stress conditions (temperature and food availability) [8, 47]. Similarly, expression of key biomineralization genes such as *nacrein* or *pif* in pearl sacs was clearly dependent on the time of sampling post-graft [8, 23]. *Pif*-related genes are particularly interesting because they directly controlled crystal growth during an *in vitro* experiment [41], yet the clear connexion between *pif*-related gene expression and pearl phenotypes remains unclear. The present study showed that neither *pif* nor *pif-177* were differentially expressed between high and low pearl quality despite these genes having already been correlated with pearl weight (Rho = 0.259; *p*-value = 0.01) and pearl quality 3 months post-graft in previous studies on *P. margaritifera* [23, 48]. Inversely, the *BMSP-like* gene, a gene related to the *pif* family [49], was up-regulated in high quality pearls. Clearly, further studies will be needed to unravel the complex role of *pif*-related genes during the different stages of pearl formation. Nevertheless, this study supports previous findings on the role of *aspein* and *shematrin 9* genes in controlling pearl quality, independently of geographical location or time of post-graft sampling. Inversely, *pif* gene (*pif-177, pif* and *BMPS-like*) expression was not consistent with previous studies and suggests that *pif* expression variation is strongly time-dependent, which could be the basis of the complex role of the *pif*-related gene family in pearl biomineralization.

### Different exon usage plays a role in shaping pearl quality

Genes and related proteins involved in biomineralization have complex structures and often require post-translational modifications and proteolytic cleavages [50]. Alternative splicing in biomineralization-related genes has already been suggested by the presence of numerous isoforms for spicule matrix protein (SpSMs) coding gene in sea urchin [51]. In the present study, four biomineralization-related genes, *pif, aspein, perlwapin* and *wfd18*, were found to have significant differences in exon usage, and were all also differentially expressed between low and high pearl quality. The present study could not, however, assess whether the control of exon usage is under epigenetic and/or genetic control. In an effort to reduce the variability inherent to the complex determination of pearl quality, our sampling design only included pooled samples from a single geographic location and mixed several pearl defect types. However, broadening our results on splicing events to the individual level and specific defect types could enable us to redefine the link between biomineralization gene architecture and pearl quality traits.

### Transposable elements might be involved in the regulation of cultured pearl quality

Strikingly, this study only identified a few genes as being up-regulated in high quality pearls (n = 16) compared to the number of up-regulated genes in low quality pearls (n = 246). Among these up-regulated genes, there was a surprisingly high representation of transposable element- (TE) related genes (n = 3 out of 7 annotated genes) including long-transposable elements (LTR) of the Gypsy and Jockey families. Recent studies in humans and plants suggest that TEs and TEs insertion are effective regulators of gene expression and alternative splicing events [52–54]. Furthermore, it has been shown that both tandem repeats (TRs) and TEs are intimately linked with TRs derived from younger/more active TEs [55]. As an example, a study in maize supports the results of specific centromeric TRs originating from Ty3/gypsy retrotransposons [56]. From an evolutionary perspective, genes involved in biomineralization are structurally complex and often characterized by the presence of numerous TRs [57]. It is thus plausible that control of tandem repeat formations might result from TE insertion [44, 45]. However, by which mechanisms TEs (specifically Gypsy and Jockey family TEs) are involved in pearl quality remains to be elucidated. Further studies correlating TE insertion with the structure of transcripts (such as *aspein*), considering the specific genotypic background, should provide useful information that will help unravel the complex regulation of cultured pearl quality traits.

## Conclusions

This study successfully made it possible to: 1) identify genes whose expression in pearl sacs was associated with cultured pearl quality in *P. margaritifera*, and 2) highlight other putative regulation levels for pearl quality determination through alternative splicing and TE regulation. Among the genes differentially expressed, new candidates were identified for pearl quality (*perlwapin, BMSP-like*), as were previously described biomarkers (*aspein* and *shematrin-9*). The present study also showed, however, that gene expression of some biomarkers previously associated with pearl quality or thickness (*pif-177, pif, msi30, pearlin* or *nacrein*), is not systematically correlated with pearl quality, suggesting that other factors might be involved. Further studies should focus on time-course experiments from the first stages of mineral deposition until harvest so as to unravel gene expression in the successive biomineralization events and the interplay between environmental and genetic factors in controlling specific quality traits.

## Methods

### Transcriptome reference assembly of *P. margaritifera*

The multi-tissue reference transcriptome was built with tissues obtained from a total of 10 *P. margaritifera* individuals: gonad tissues (n = 2, obtained from one male and one female; [36]), whole tissue of 4-month-old juveniles (whole individuals, n = 2) and mantle tissue (n = 6; Table 1). For all samples apart from the gonads, total RNA was extracted with TRIZOL Reagent (Life Technologies) at a ratio of 1 ml per 100 mg tissue, following manufacturer’s recommendations. RNA quantity/integrity and purity were validated on a Nanodrop (NadoDrop Technologies Inc.) and on a BioAnalyzer 2100 (Agilent Technologies), respectively. RNA was dried in RNA-stable solution (ThermoFisher Scientific) following manufacturer’s recommendations and shipped at room temperature to McGill sequencing platform services (Montreal, Canada). TruSeq Sample Prep. (Illumina, San Diego, Ca, USA) RNA libraries were multiplexed (n = 10 pools by lane) and sequenced by HiSeq4000 100-bp paired-end (PE) sequencing technology. For the gonads, the samples were also sequenced by HiSeq2000 100-bp PE and were downloaded from the SRA database (Bioproject PRJNA229186; see [36] for more information on RNA preparation). Reads were filtered for adapter removal, minimum length (≥ 40-bp) and minimum quality (Q = 28) using Trimmomatic v0.36 [58]. The retained PE reads were assembled with Trinity v2.4.0 [59] using default parameters with a minimum transcript length of 200-bp. Read quality was assessed before and after read trimming with FastQC v0.11.5 (https://www.bioinformatics.babraham.ac.uk/projects/fastqc/).

**Table 1:**
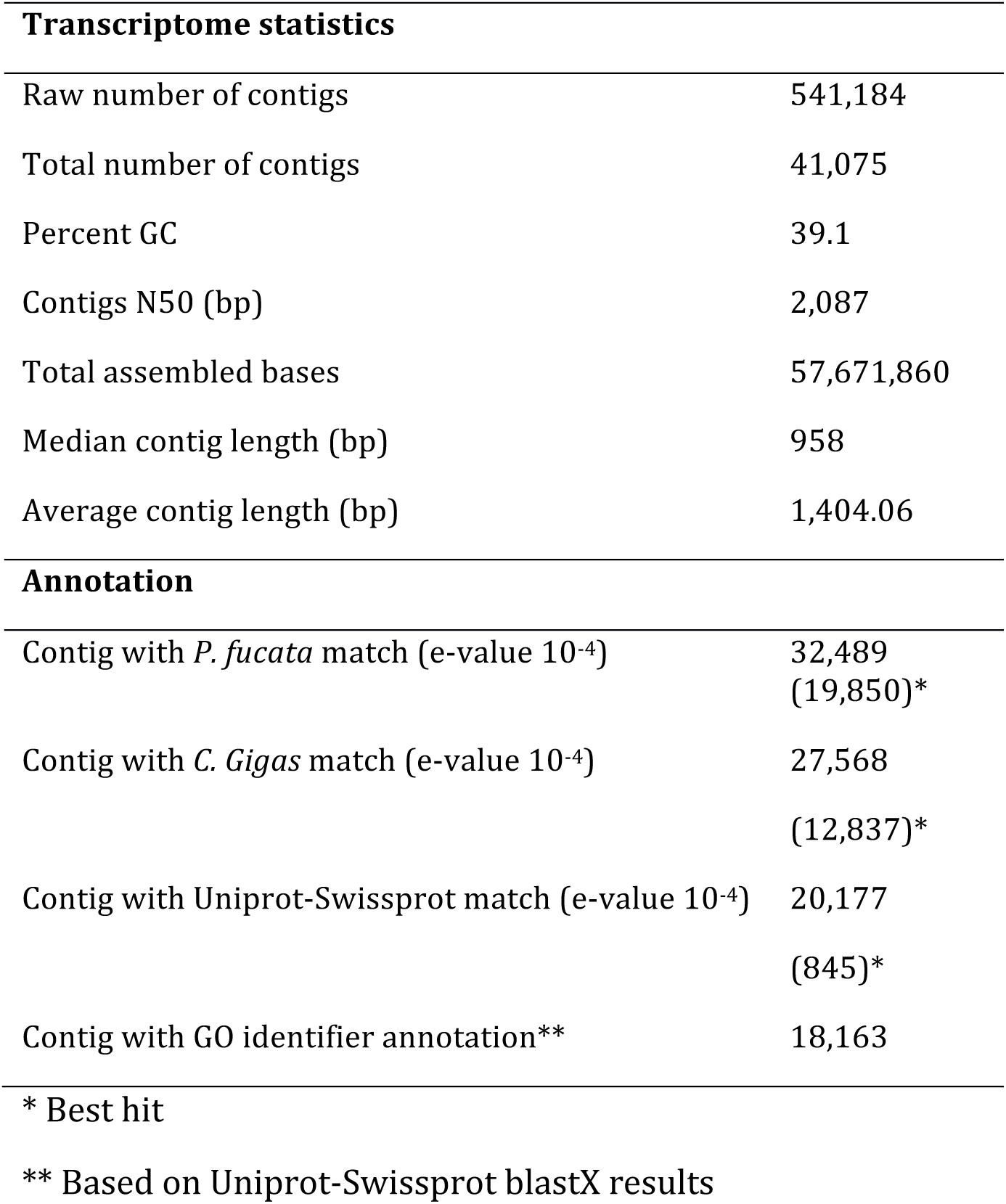
Transcriptome assembly, annotation statistics and differential expression results

Functional and mapping-rate filtering approaches were combined to reduce the redundancy present in the first version of the *P. margaritifera* transcriptome. Briefly, open-reading frames (ORFs) for each transcript were predicted using ‘*LongOrfs*’ function implemented in Transdecoder v3.0.1 [59]. Only the transcripts containing an ORF of at least 100 amino acids were kept. Another filtering step included the removal of isoforms with residual expression; hence, only the most expressed isoforms for each gene with a mean mapping rate of 0.5 transcripts per million (TPM) were kept. We used a blastN approach against curated and non-redundant viral, bacterial, archeal, plasmid and fungal RefSeq transcripts databases (Download 19-09-17; ftp://ftp.ncbi.nlm.nih.gov/refseq/release/) to remove putative contamination. Transcripts matching a reference with an e-value < 10^-10^, minimum identity of 75% and minimum query coverage of 70% were discarded. Finally, Illumina adapters were screened in the transcriptome using a blastN approach and adapter list (http://omicsoft.com/downloads/ngs/contamination_list/v1.txt). Assessment of the final transcriptome completeness was conducted with BUSCO v1.1b [60] against the conserved single-copy metazoan genes database (n = 978). Each filtering step was validated with TransRate v1.0.3 [61]. Filtered reads were then mapped back on the filtered reference transcriptome to evaluate individual mapping rate with GSNAP v2017-03-17 [62].

For functional annotation, the transcripts were searched against Uniprot-Swissprot [63], *Pinctada fucata* (http://marinegenomics.oist.jp/pearl/viewer/download?project_id=20) and *Crassostrea gigas* (ftp://ftp.ncbi.nih.gov/genomes/Crassostrea_gigas/RNA) protein databases using a blastX approach (e-value < 10^-4^) [64]. The best hit results are reported in Table 2.

### Animal and tissue sampling

An experimental graft was realised in order to obtain cultured pearls and their corresponding pearl sacs. For this, a total of 20 pearl oyster donors were used to perform 600 grafts (30 grafts per donor) over a 2-day period, using 2.4 BU nuclei (7.304 mm diameter, 0.59 g weight -Nucleus Bio, Hyakusyo Co., Japan) in December 2015. All grafts were performed under standard production conditions by a single expert at the Pahai Poe Pearl Farm (Apataki atoll, 15°34’S, 146°24′ W, Tuamotu archipelago, French Polynesia) so as to minimize the grafter effect on pearl quality traits described in [65]. *Pinctada margaritifera* used in this part of the study had been collected as wild spat in December 2013 using commercial collectors in the lagoon of Takapoto atoll (14°32′ S, 145°14′ W, Tuamotu archipelago, French Polynesia), two years before the experimental graft took place.

At the time of pearl harvest (18 months after the grafting operation, May 2017) and in order to minimize the contamination by recipient tissues, the gonad was first cut from the recipient oysters. The gonad tissue was then removed with a surgical blade until only a thin (< 0.5 mm) layer of tissue surrounding the pearl remained. At this point, only the pearl sac and the pearl remained [8]. Next, an incision was made in the pearl sac in such a way as to remove the pearl, which was then placed in a numbered box for traceability. The pearl sac was kept in a 2.0 ml tube with RNAlater until RNA extraction. A total of 442 pearl sacs were sampled (73.6% of the total number of oysters grafted). This pearl harvest rate represents the production yield, as nucleus rejection and mortality were observed over the entire culture time [66].

Cultured pearl grade was evaluated as described in [67]. The pearls were cleaned by ultrasonication in soapy water (hand washing) with a LEO 801 laboratory cleaner (2 L capacity, 80 W, 46 kHz); they were then rinsed in distilled water. To ensure homogeneity in parameter assessment, all evaluations were made visually (no magnification devices were used) by the same professional operator. Cultured pearl grade is made up of two components: surface defects and lustre. When pearls are graded, the appearance of their surface is evaluated. The degree of imperfection is correlated with the number of defects (i.e. smooth surface *vs.* one or more spots). For the grade classification, high pearl quality corresponds to grades of C and above, and low quality to D1 and D2 grades (Figure 1). From the high and low quality pearls, twenty pearl sacs from each group that had produced only green pearls were randomly selected. Overall, high quality pearls were significantly heavier, with a thicker nacre layer than the low quality pearls (t-test; *p*-value < 0.05). Our sampling included 10 donors equally allocated between several pools (either low or high pearl quality, as illustrated in Figure 1) with the objectives of reducing putative donor effect [68–71].

### RNA extraction and sequencing

The total RNA extraction procedure was identical to that for the samples used in the transcriptome assembly. For each condition (low or high pearl quality), an equimolar RNA quantity was pooled (n = 5 individuals/pool) to give a total of five pools by condition. RNA was dried in RNA-stable solution (ThermoFisher Scientific) following manufacturer’s recommendations and shipped at room temperature to McGill sequencing platform services (Montreal, Canada). TruSeq Sample Prep. (Illumina, San Diego, Ca, USA) RNA libraries were multiplexed (N=10 pools by lane) and sequenced by HiSeq 4000 100-bp paired-end (PE) sequencing technology.

### Differential expression analysis

Raw reads were trimmed using Trimmomatic v0.36 [58], using similar parameters as for the transcriptome assembly. Only paired-end reads were retained and mapped to the reference transcriptome, using GSNAP v2017-03-17 [62] with default parameters but allowing a minimum mismatch value of 5. Low mapping quality, multi-mapping (-q 5) and “non-properly paired reads” (-f 0×2) were removed using Samtools v1.4.1 [72]. A matrix of raw counts was built using HTSeq-count [73]. Low coverage transcripts (CPM < 1 in at least 10 individuals) were removed, resulting in a total of 40,633 transcripts and differential expression was assessed using the DESeq2 R package [74]. Transcripts were considered significant when FDR < 0.01 and |logFC| ≥ 1. Gene ontology (GO) enrichment was tested using GOAtools v0.6.5 [75] and the go-basic.obo database (release 2017-04-14) using Fisher’s test. Our background list included the ensemble of genes used for differential expression after filtering for low coverage (n = 40,633 transcripts). Only GO terms with *p*-value < 0.05 and including at least three differentially expressed genes were considered. Significant GO enriched terms were used for semantic-based clustering in REVIGO (http://revigo.irb.hr/).

### Individual gene expression validation

RT-qPCR was used to validate the expression patterns in the pearl sac, observed in RNA-seq, for key genes commonly used as markers of pearl quality traits, namely *pif-177, aspein, shematrin-9* and the *mantle protein 10* (MP10). For each pool, total RNA was treated with DNAse I using a DNA-free Kit (Ambion). First, strand cDNA was synthesized from 500 ng total RNA using the Transcriptor First Strand cDNA Synthesis Kit (Roche) and a mix of poly (dT) and random hexamer primers. Real-Time PCR amplifications were carried out on a Roche Light Cycler 480. The amplification reaction contained 5 μL LC 480 SYBR Green I Mast (Roche), 4 μL cDNA templates, and 1 μL of primer (1μM), in a final volume of 10 μL. Each run included a positive cDNA and a blank control for each primer pair. The run protocol was as follows: initial denaturation at 95°C for 10 min followed by 40 cycles of denaturation at 95°C for 30 s, annealing at 60°C for 30 s and extension at 72°C for 60 s. Lastly, the amplicon melting temperature curve was analysed using a melting curve program: 45–95°C with a heating rate of 0.1°C s^-1^ and continuous fluorescence measurement.

All measurements were made in duplicate and all analyses were based on the Ct values of the PCR products. Relative gene expression levels were calculated using the delta–delta method, normalized with two reference genes (*elongation-factor 1* and *glyceraldehyde-3-phosphate dehydrogenase*), to compare the relative expression results as follows: 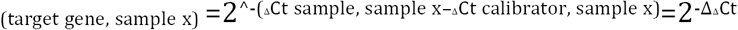 [76]. Here, the ΔCt calibrator represents the mean of the ΔCt values obtained for the tested gene. The delta threshold cycle (ΔCt) is calculated as the difference in Ct for the target and reference genes. The relative stability of the GAPDH and EF-1 combination was confirmed using NormFinder [77]. PCR efficiency (E) was estimated for each primer pair by determining the slopes of standard curves obtained from serial dilution analysis of a cDNA to ensure that E ranged from 90 to 110%. The primers used for amplification are listed in Table S2. Wilcoxon non-parametric tests were used to compare relative expression between conditions, differences were considered significant when p-value < 0.01. This complete list of primers is given in the supplementary file (Table S3).

### Alternative gene splicing and exon usage

To detect putative differential splicing variants, the reads were mapped on the scaffolds of the assembled draft genome, available for *P. margaritifera*. To reduce non-biological redundancy inherent to the current assembly state of the genome, only scaffolds with length >3,000 bp were used for the mapping. From the 757,552 scaffolds initial set, only the 32,705 longest scaffolds were retained for downstream analysis, on which 37,662 (92.7%) of the transcripts in our set could be positioned using the GMAP v2017-03-17 aligner with the default parameters for annotation [78]. Reads were mapped on the filtered reference genome using GSNAP v2017-03-17 aligner allowing five mismatches, splicing and using the ‘*splitting-output*’ function to retain only concordant and unique mapped paired-end reads [78]. We used the QORTs [79] and JunctionSeq R packages [80] to detect significant differences in exon usage. Only exons and junctions with a minimal coverage of six were used for the analysis and only differences with FDR < 0.01 were considered significant.

## List of abbreviations

BMSP-like: Blue mussel shell protein-like
Pwap: Perlwapin
wfd18: Wap four-disulfide core domain protein

## Declarations

### Availability of data and material

Raw transcriptome sequences data have been deposited at NCBI (Bioproject: PRJNA449941). Genome and transcriptome assemblies will be made publicaly available upon acceptance of the manuscript. Codes for RNA-seq analysis are available upon request.

### Authors’ contributions

CLK conceived the study. NL, VQ conducted the pearl sac sampling and the RT-qPCR analysis, respectively. PA and JLL assembled the transcriptome. CR and JVD assembled the reference genome of *P. margaritifera*. JLL analysed the RNAseq data. JLL and CLK wrote the paper. All co-authors contributed to reviewing the manuscript and accepted the final version for publication.

## Acknowledgements

The authors would especially like thank the host site, Pahai Poe Pearl Farm located in Apataki atoll (Tuamotu archipelago, French Polynesia) for their generous support, and D. Potin for his assistance with the cultured pearl photographs. The authors thank the Ifremer (Datarmor; RIC) and Station Biologique de Roscoff (ABiMS; http://abims.sb-roscoff.fr) bioinformatic platforms for providing computational resources and analysis support.

## Supplementary

Figure S1: Summarized REVIGO semantic plot for gene ontology enrichment analysis

Figure S2: Relative gene expression for biomineralization genes analysed by qPCR in the pearl sac of *P. margaritifera*. Values are expressed as means of relative expression ± standard deviation. Asterisks indicate significant differences (Wilcoxon test, *p*-value < 0.01).

Table S1: *P. margaritifera* individuals used for the transcriptome assembly. NA = Not identified.

Table S 2: Complete list and statistics on differentially expressed genes and their annotation.

Table S 3: Set of forward and reverse primers used for the biomineralization gene expression (real-time PCR) analysis in *Pinctada margaritifera*.

